# Ovarian Hormones and Obesity Drive Th17-mediated Airway Inflammation through Estrogen Receptor Signaling

**DOI:** 10.64898/2025.12.02.691857

**Authors:** Emely Henriquez-Pilier, Jacqueline-Yvonne Cephus, Shelby Kuehnle, Elie Tannous, Alessandra Tomasello, Kaitlin McKernan, R. Stokes Peebles, Katherine N. Cahill, Jeffrey C. Rathmell, Dawn C. Newcomb

## Abstract

Obesity is a risk factor for increased prevalence and severity of asthma, particularly in females. As adults, females have increased prevalence of asthma compared to males. Yet, the mechanisms remain unclear on how sex hormones and obesity increase airway inflammation. We hypothesize that estrogen signaling through estrogen receptor-alpha (ER-α) in T cells increased airway inflammation in the context of obesity. To test our hypothesis, we utilized a high fat (HFD) on female and male mice that underwent ovariectomy or gonadectomy or in Esr1^fl/fl^ X Cd4^Cre+^ male and female mice. As controls, mice in similar groups were fed normal chow. After 8-12 weeks on diets, house dust mite (HDM) sensitization and challenge occurred in all mice. Lungs and BAL fluid were harvested 24 hours after the last challenge. Ovarian hormones and ER-α signaling in T cells increased eosinophils, neutrophils, and Th17-mediated airway inflammation in the lungs of obese female mice. Additionally, using PBMCs from a well-characterized obese asthma cohort, we determined that obese women with asthma had increased Th17 cells compared to obese men with asthma. Our results show that ER-α signaling in T cells increases Th17-mediated airway inflammation in obese mice and that Th17 cells circulate at higher frequencies in women with asthma compared to men with asthma. Further research into the interplay between hormonal signaling and immune responses in asthma is essential for developing personalized treatments.

**One Sentence Summary:** Estrogen receptor-alpha signaling, in the context of obesity, increases allergen-induced Th17-mediated airway inflammation in female mice.

## Introduction

The current obesity epidemic has established obesity as a significant risk factor for severe, uncontrolled asthma(1–3). Among adults, obese females demonstrate a markedly higher prevalence of asthma compared to both non-obese females and males across all weight categories(1, 2, 4, 5). Yet, the mechanisms driving this heightened asthma susceptibility in obese females remains unclear.

Airway inflammation in asthma is characterized by type 2 and non-type 2 inflammation. Type 2 inflammation results in increased IL-4, IL-5, and IL-13 production from CD4^+^ Th2 cells and ILC2 leading to increased eosinophil infiltration, IgE production, airway hyperresponsiveness (AHR), and mucus production. Non-type 2 inflammation leads to increased IFN-у and IL-17 from CD4^+^ Th1, Th17, ILCs, or NK cells, resulting in increased neutrophilic inflammation, AHR, and mucus production. High fat diet (HFD)-induced obesity in mice results in increased type 2 and non-type 2 airway inflammation, including neutrophil infiltration, compared to non-obese mice(6–8).

Previous studies in our laboratory showed that sex hormones modify airway inflammation in non-obese mice. Ovarian hormones increased allergen-induced airway inflammation while androgens decreased allergen-induced airway inflammation(9–15). Ovariectomy (OVX) in female mice, which reduced the production of ovarian hormones, attenuated allergen-induced eosinophil and neutrophil infiltration into the airway and AHR compared to sham-operated female mice, with normal levels of ovarian hormones(9). Estrogen and progesterone were also required for Th17 cell differentiation in both mice and humans(9). Additional studies showed that estrogen signaling through estrogen receptor alpha (ER-α), but not estrogen receptor beta (ER-β), increased Th17 cell differentiation and IL-33-mediated type 2 inflammation(10, 11). Additionally, females deficient in ER-α (*Esr1*^−/-^ mice) exhibited markedly reduced airway inflammation with diminished eosinophil and neutrophil infiltration compared to WT females. Yet, how sex hormones altered airway inflammation in the context of obesity remained unclear. We hypothesized that ER-α signaling in CD4⁺ T cells enhance Th2- and Th17-mediated airway inflammation in obese female mice. Using a high-fat diet (HFD) model to induce obesity, we determined that ER-α signaling in T cells increased Th17-mediated airway inflammation in obese female mice. These results were then confirmed in PBMCs from obese women with moderate-to-severe persistent asthma showing increased numbers of Th17 cells compared to obese men with asthma.

## Methods

### Mice

All experiments were performed at Vanderbilt University in accordance with Institutional Animal Care and Utilization Committee (IACUC)-approved protocols and conformed to all relevant regulatory standards. Mice were housed in a pathogen-free facility with five mice per cage and ad libitum access to food and water. Sham-operated female and male mice, ovariectomized (OVX) female mice, and gonadectomized (GNX) male mice were obtained from Jackson Laboratories and surgeries were performed at 3-4 weeks of age at Jackson Laboratories.

*Esr1^fl/fl^* (strain 032173) and *Cd4^Cre^* (strain 022071) were purchased from Jackson Laboratories and then crossed to generate *Esr1^fl/f^* X *Cd4^Cre^* in our laboratory. Mice were genotyped using Transnetyx. 6 to 8-week-old male and female mice were used for all animal experiments.

### Diet

45 kcal% HFD was obtained from Research Diets (cat. D12451) and used for HFD groups. Control diet groups used LabDiet’s 4.5 kcal% fat diet (cat. 5LOD). Treatment groups were placed on their respective diets for 8-12 weeks with body weight measured every other week.

### House dust mite (HDM) allergen exposure

After 8-12 weeks of HFD or control diet, mice underwent house dust mice (HDM) allergen sensitization and challenge. Mice were anesthetized with isoflurane and challenged intranasally sensitized with 25 µg of HDM in 80µl of PBS on days 0, 3. Mice were then challenged intranasally with 25 µg of HDM on days 14, 15, 16 and 17. Twenty-four hours after the last challenge, mice were sacrificed for collection of lungs and downstream analysis.

### BAL

After sacrifice, BAL fluid was obtained through the insertion of a tracheostomy tube, instillation of 800 µL of PBS through a syringe into the lungs, gentle massage of the lungs, and gentle withdrawal of fluid through the same syringe. Total cells were determined, and BAL cell suspension were adhered to a slide and stained using a Three-Step Stain set from Epredia Ref: 3300. Using light microscopy, BAL cells were classified as eosinophils, neutrophils, macrophages/monocytes or lymphocytes. The percentage of each cell type was determined from the total number.

### Flow Cytometry

Lungs were harvested, minced, and digested using 1 mg/mL collagenase type and DNase I in RPMI with 10% FBS for 30 minutes and 37°C. Digestion was stopped using 1 µM EDTA. Cells were filtered through a 70 µm strainer to remove debris and red blood cell lysis was performed. Cells were restimulated in IMDM with GlutaMax and HEPES, 10% FBS, 1% penicillin/streptomycin, 1% sodium pyruvate, 50 µM 2-mercaptoethanol, 1 µM ionomycin, 50 ng/mL PMA, and 0.07% Golgi Stop at 37°C for 4 hours. Cells were then washed with PBS and 3 million viable cells were used for flow staining. Cells were stained with a fixable viability dye (Live Dead Aqua) and blocked using an anti-mouse FcR antibody at 4°C. Cells were washed with PBS with 3% FBS and stained for surface markers (Supplementary Table 1) for 45 minutes at 4°C. Cells were again washed with PBS with 3% FBS and were subsequently fixed and permeabilized using the FoxP3 Transcription Factor Fix/Perm Kit for unstimulated cells and Cytofix/Cytoperm Kit for restimulated cells. After washing with the provided Perm Wash, cells were stained for intracellular markers (Supplementary Table 1) as stated in the text for 45 minutes at 4°C. Cells were washed again with Perm Wash and once more with PBS with 3% FBS. Flow cytometry was conducted on a Cytek Aurora and data were analyzed using FlowJo v.10.

### Human PBMC samples recruitment, collection, and assessment

Human PBMCs and patient demographics were collected and obtained at baseline from participants in the Glucagon-like Peptide-1 Receptor Agonist in the Treatment of adult, obesity-related, symptomatic Asthma clinical trial (GATA-3, clinical trial #NCT05254314). Participants eligible for inclusion in our analysis were adults (18-50 years), capable of providing informed consent, with physician-diagnosed persistent asthma requiring at least medium-dose inhaled corticosteroids or more therapy, symptomatic disease (Asthma control questionnaire (ACQ)-6 score ≥1.5), and evidence of bronchodilator responsiveness or airway hyperresponsiveness. All participants had either a BMI ≥30 kg/m^2^ or BMI≥27 kg/m^2^ with at least one weight-related comorbidity. Female participants of childbearing potential must test negative for pregnancy. Key exclusion criteria were, asthma biologic therapy use, diabetes mellitus (HbA1c >6.5), excessive use of short-acting bronchodilators, oxygen saturation <94% on room air, recent tobacco, e-cigarette, or marijuana use, and use of exogenous sex hormones (including oral birth control medications).

PBMCs were plated at 75,000 cells per well in a 96-well non tissue culture–treated U bottom plate. T cells were activated overnight at 37°C with 5% CO_2_ with Dynabeads Human T-activator CD3/CD28/CD137 (Thermo Fisher Scientific Ref: 11163D) and rhIL-2, in warmed T cell media (500mL HPLM, 10% FBS, RPMI 1640 100X Vitamins, 1% Pen/Strep). After overnight stimulation, cells were refreshed with warmed T cell media and allowed to equilibrate for 1 hour. Cells were then washed with PBS and stained with Live Dead Far Red fixable viability dye for 20 minutes at 4°C. Subsequently, cells were washed with PBS with 3% FBS and stained at 37°C with 5% CO2 for surface chemokine markers for 20 min and then the remainder of surface markers were applied at room temp for 30min (Supplementary Table 2). Cells were washed again with PBS with 3% FBS and fixed using the Foxp3 Transcription Factor Fix/Perm Kit (ThermoFisher Scientific) per manufacturer instructions and intracellular staining and flow cytometry was performed as described above.

### Statistical analysis

Statistical analyses were performed using GraphPad Prism (v. 10) with data represented as mean ± SEM or mean ± standard deviation as denoted in the legend. Data were significant when *p* was less than 0.05. Comparisons between 2 groups were performed using 2-way *t* test or Mann-Whiteny *U* test for parametric or nonparametric data, respectively. Comparisons of greater-than 2 groups were made using an ANOVA with post hoc for multiple comparisons.

## Results

### Ovarian hormones and HFD-induced obesity increased HDM airway inflammation

To understand if obesity and ovarian hormones increased allergic airway inflammation, WT female mice underwent an OVX at 3-4 weeks of age to remove the ovaries and limit ovarian hormones. Male mice underwent a GNX at 3-4 weeks of age to remove the testes and limit androgen production. As a control, WT female and male mice also underwent sham operations (Sham). At 6-8 weeks of age, mice were placed on a HFD that contained 45% fat or normal chow (control diet) (4.5% fat) for 12 weeks as shown in Figure 1A. While on HFD or normal chow diets, mice were weighed every 2 weeks. Sham operated female mice on control diet and HFD had similar weight gain over the 12 weeks (Figure 1B). OVX female mice on control diet had increased weight gain compared to sham-operated female mice (on any diet?), and OVX female mice on HFD exhibited (can we just say: the greatest weight gain?) more weight gain than OVX female mice on control diet or all sham-operated female mice. Sham-operated and GNX male mice on HFD had increased, but similar, weight gain compared to sham-operated and GNX male mice (Figure 1B). After 12 weeks on HFD or normal chow, fasting glucose levels were determined and male and female mice on HFD had increased levels of fasted serum glucose compared to male and female mice fed normal chow (Supplemental Figure 1).

**Figure 1:**
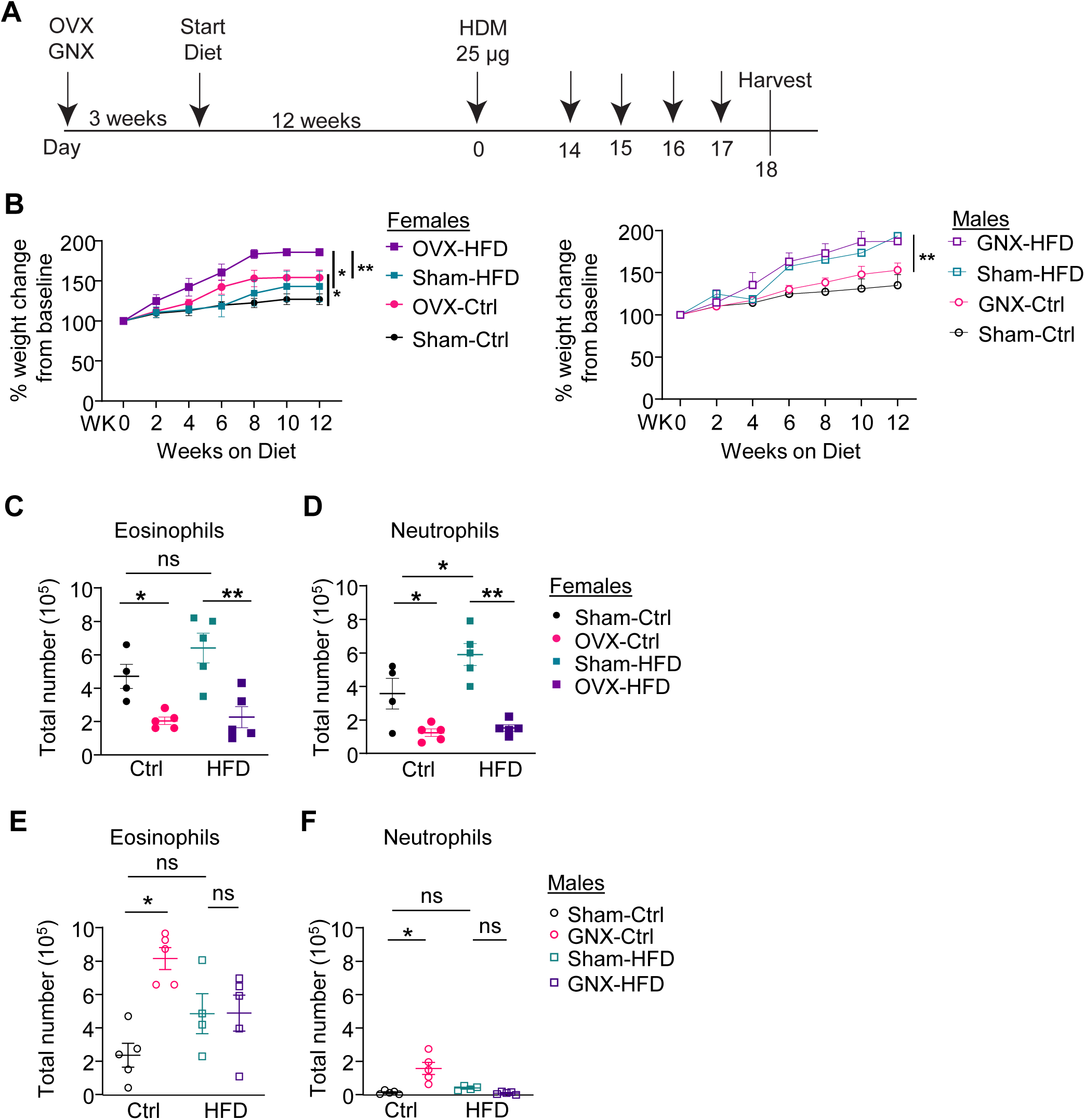
Ovarian hormones and HFD increase HDM-induced airway inflammation. **A)** Timeline showing experimental protocol. Briefly, mice underwent sham operation, ovariectomy (OVX) for females, or gonadectomy (GNX) for males at 3-4 weeks of age. Three weeks following the surgeries, mice were administered regular chow (control) or high-fat diet (HFD) for 12 weeks. Mice were then intranasally sensitized and challenged with 25µg of house dust mite (HDM) with diet regiment maintained. Lungs and BAL fluid were harvested 24 hours following last HDM challenge. **B)** Percent change in body weight from baseline in female (left) and male (right) mice fed HFD or control diet. Data are expressed as mean ± SD; n = 4-5 mice per group, *p < 0.05, **p < 0.01 two-way ANOVA with Tukey’s post-hoc test. **C–D)** Quantification of BAL eosinophils, neutrophils in females. **E-F)** Quantification of BAL eosinophils, neutrophils in males. C-F) Data are expressed as mean ± SEM; n = 4-5 mice per group; *p < 0.05, **p < 0.01 one-way ANOVA with Bonferroni post-hoc test.

All mice were then sensitized and challenged with HDM as depicted in Figure 1A. Twenty-four hours following the last challenge, lung tissues and bronchoalveolar lavage (BAL) fluid were collected. Analysis of BAL fluid cell counts revealed that sham-operated female mice on the control diet showed increased eosinophils and neutrophils in BAL fluid compared to the OVX female mice on control diet (Figure 1C-D). With HFD, sham-operated female mice had increased BAL eosinophils and neutrophils compared to OVX female mice. Additionally, HFD fed, sham-operated female mice had increased BAL neutrophils compared to sham-operated females on the control diet (Figure 1D). In the male mice, GNX male mice on control diet had increased eosinophils and neutrophils compared to sham-operated male mice on control diet (Figure 1E-F). No differences were detected in sham-operated and GNX male mice on HFD.

We next examined whether sex hormones influence the IL-13 and IL-17A-producing CD4^+^ T cells in the lungs. IL-13 and IL-17A producing CD3^+^ CD4^+^ cells were determined by flow cytometry using the gating strategy shown in Figure 2A, and Supplemental Figure 2. IL-13^+^ CD3^+^ CD4^+^ T cells were denoted as Th2 cells, and IL-17A^+^ CD3^+^ CD4^+^ T cells were denoted as Th17 cells. HFD fed sham-operated female mice had increased lung Th2 and Th17 cells compared to control diet sham-operated female mice and HFD OVX female mice (Figure 2B-C). GNX male mice on control diet had increased numbers of lung Th17 cells compared to sham-operated male mice on control diet (Figure 2E), but no differences in Th2 or Th17 cells in sham-operated and GNX male mice on HFD were detected (Figure 2D-E). Overall, these data shows that ovarian hormones and HFD diet increased HDM-induced airway inflammation.

**Figure 2:**
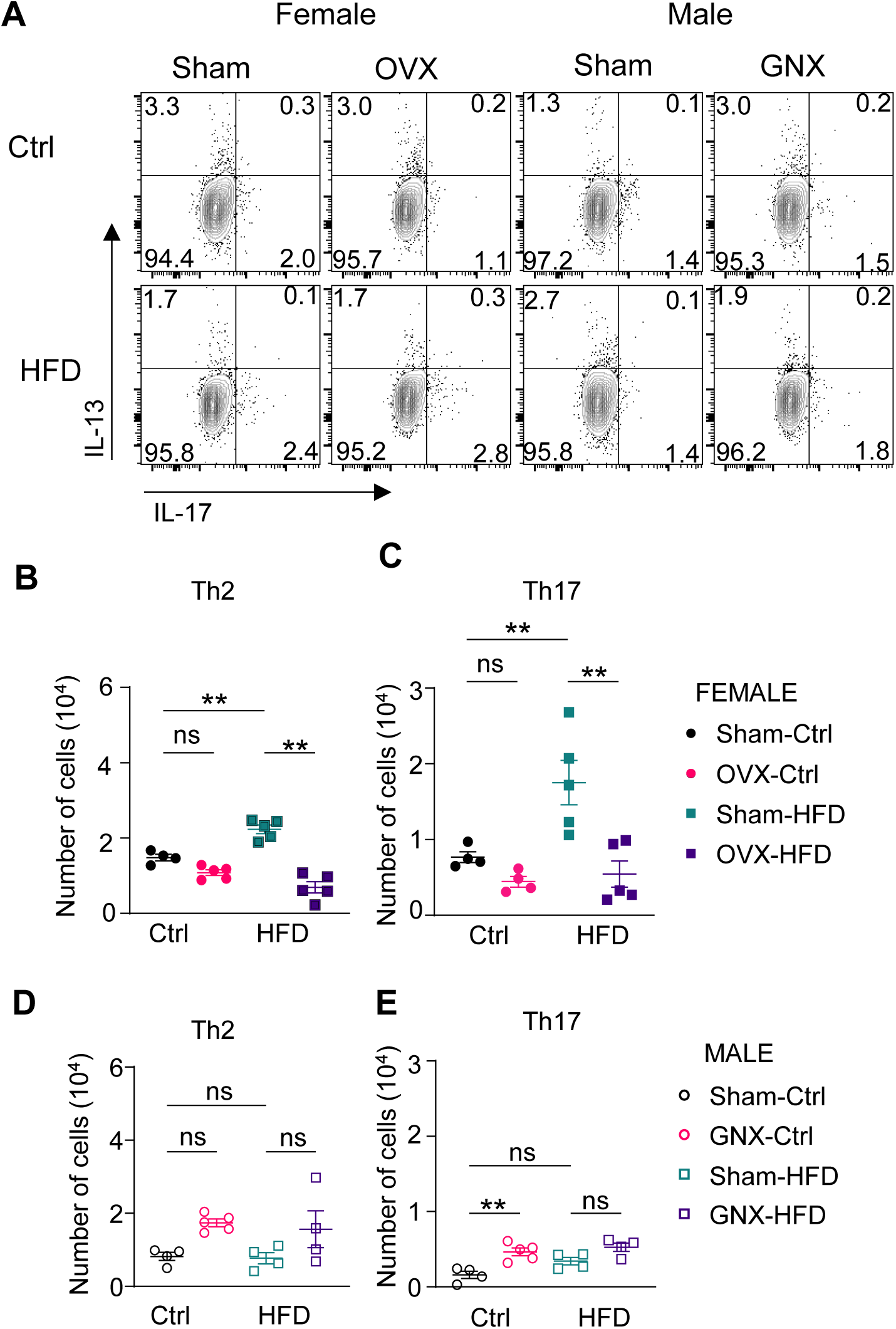
Ovarian hormones and HFD increase HDM-induced lung Th2 and Th17 cells. Lungs were harvested 24 hours following the last HDM challenge and assessed for CD4+ Th2 and Th17 cells. **A)** Representative dot plots of IL-13+ Th2 cells and IL-17A+ Th17 cells in HDM-challenged female and male mice on either control diet (Crtl) or HFD. Cells were pre-gated on viable, CD45+, CD3+ CD4+ cells. **B-E)** Quantification of lung Th2 and Th17 cells in female mice (panels B-C) and male mice (panels D-E). Data are expressed as mean ± SEM; n = 4-5 mice per group, **p < 0.01 one-way ANOVA with Tukey’s post-hoc test.

### Estrogen receptor-α signaling in CD4^+^ T cells increased HFD and HDM-indued airway inflammation

To determine whether estrogen receptor-alpha (ER-α) signaling in T cells contributes to the increased airway inflammation observed in HFD female mice, we next utilized genetically modified mice lacking ER-α specifically in T cells. *Esr1*^fl/fl^ and *Esr1*^fl/fl^ *Cd4*^Cre+^ female and male mice were fed a HFD or control diet and then sensitized and challenged with HDM as described in Figure 1A. BAL fluid and lungs were harvested 24 hours after the last HDM challenge. *Esr1*^fl/fl^ female mice on HFD had increased neutrophil, but not eosinophil, infiltration in BAL fluid, compared to *Esr1*^fl/fl^ female mice on control diet (Figure 3A-B). When ER-α signaling was disrupted in T cells (*Esr1*^fl/fl^ *Cd4*^Cre+^), the HFD-induced neutrophil infiltration decreased (Figure 3B). HFD also increased Th17 cells in female *Esr1*^fl/fl^ mice, and the loss of ER-α signaling in T cells correspondingly reduced Th17 cell numbers (Figure 3E). There were no differences in neutrophil infiltration, or Th17 cells observed in *Esr1*^fl/fl^ and *Esr1*^fl/fl^ *Cd4*^Cre+^ male mice with control diet or HFD (Figure 3B, E). These results show that ER-α signaling in T cells increase HFD and HDM-induced neutrophil infiltration and numbers of Th17 cells in the lung.

**Fig. 3:**
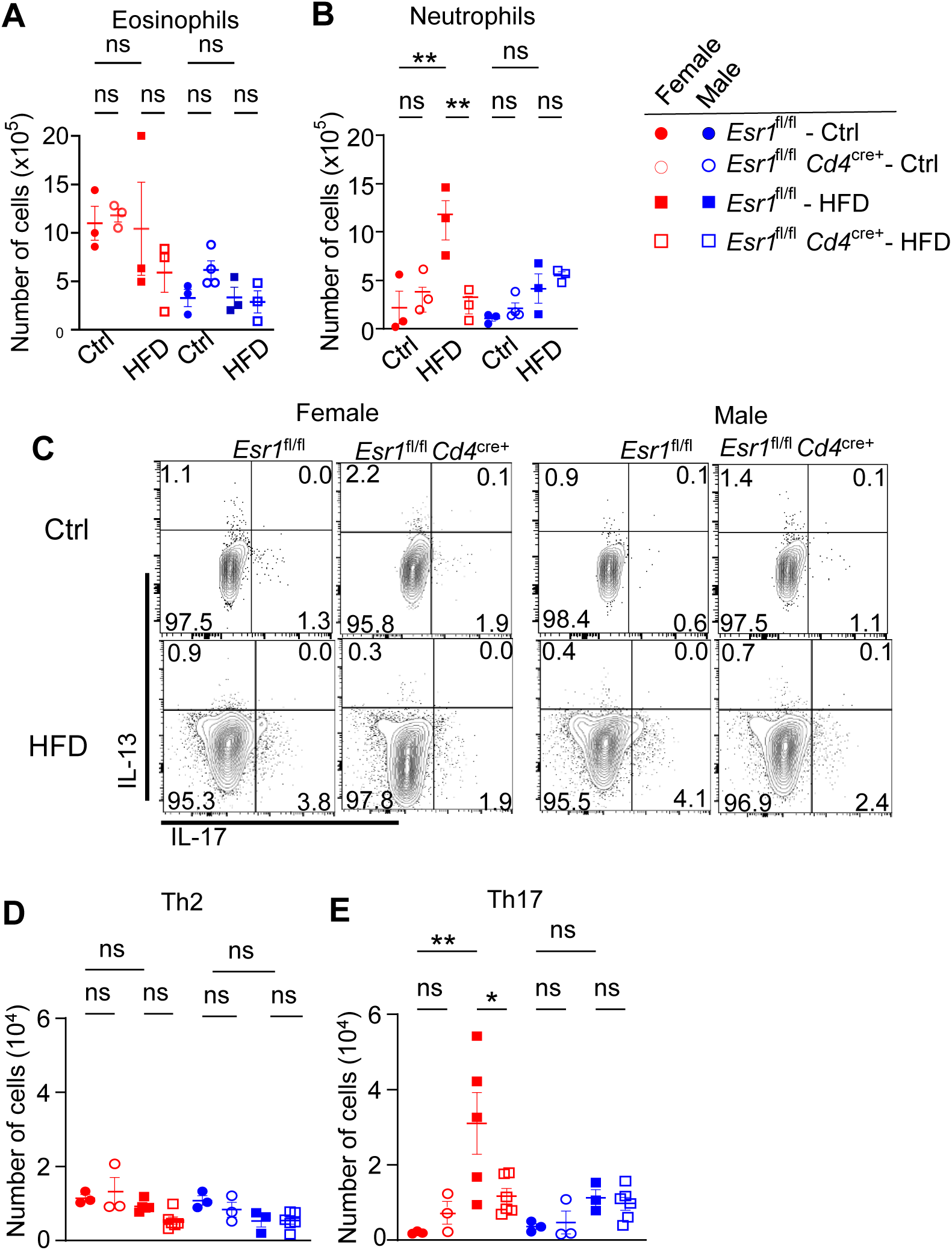
ER-α signaling in CD4+ T cells increases HFD-induced Th17 cells in HDM challenged female mice. Female *Esr1*^fl/fl^ and *Esr1*^fl/fl^ *Cd4*^cre+^ female and male mouse were put on control diet or HFD for 12 weeks followed by HDM challenge. Lungs and BAL fluid were harvested one day following the last challenge. **A-B)** Quantification of BAL eosinophils, neutrophils. **C)** Representative dot plots of IL-13+ Th2 cells and IL-17A+ Th17 cells in HDM-challenged mice on HFD. Cells were pre-gated on viable, CD3+ CD4+ cells. **D-E)** Quantification of lung Th2 and Th17 cells in female mice (panels B-C) and male mice (panels D-E). Data are expressed as mean ± SEM; n = 3-5 mice per group, *p < 0.05, **p < 0.01 one-way ANOVA with Tukey’s post-hoc test.

### Obese women with asthma have increased circulating Th17 cells compared to obese men with asthma

To investigate the influence of sex and obesity on the effector function of Th1, Th2, and Th17 cells in individuals with asthma, PBMCs were obtained from male and female participants enrolled in the GATA-3 clinical trial. Patient demographics are summarized in Supplementary Table 2. Following overnight restimulation of PBMCs with anti-CD3 and anti-CD28 to activate T cells (Figure 4A), flow cytometry was performed to assess T cell subset frequency using chemokine receptors to denote Th2 cells (CD3+ CD4+ CCR4+ CXCR3-) and Th17 cells (CD3+ CD4+ CCR6+ CCR4+/-) as shown in Supplemental Figure 4. Obese female patients with asthma had increased percentages of Th1 and Th17 cells compared to obese male patients (Figure 4B-C). Th2 cell percentages did not differ significantly between sexes. These data show that obese women with asthma have increased circulating Th17 cells compared to obese men with asthma.

**Fig. 4:**
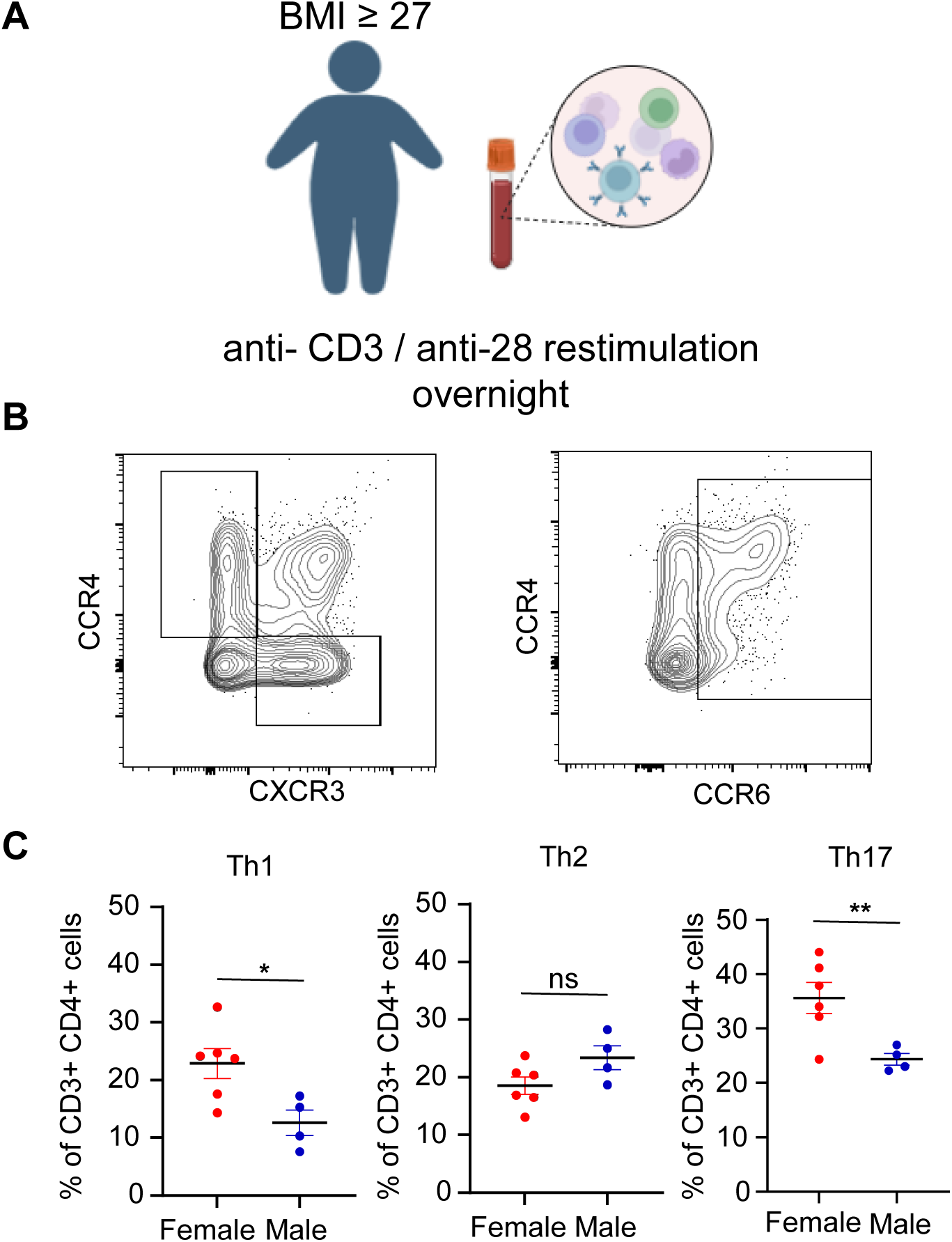
Obese women with uncontrolled asthma have increased circulating Th17 cells compared to obese men with uncontrolled asthma. **A)** Diagram of PBMC restimulation from obese women (n=6) and men (n=4) with asthma. **B)** Representative dot plot of Th1 cells (CXCR3+ CCR4-), Th2 cells (CXCR3+ CCR4+), and Th17 cells (CCR6+, CCR4+/-) from an obese women with uncontrolled asthma. Cells were pre-gated on viable, CD3+ CD4+ cells. **C)** Quantification of Th1, Th2, and Th17 percentages of CD3+ CD4+ cells. Data are expressed as mean ± SD; n = 4-6, *p < 0.05, **p < 0.01 using Welch’s unpaired T-test (*p < 0.05, **p<0.01).

## Discussion

Previous studies determined that ovarian hormones increased allergic airway inflammation in mice on normal chow(9–11). In this study, we extend these findings and show that ER-α signaling in T cells increased obesity-induced airway inflammation in a mouse model and that obese women with asthma have increased circulating Th17 cells compared to obese men with asthma. Together these findings provide mechanistic insight as to why obese women with asthma may have increased asthma prevalence and severity compared to non-obese women or all men.

Our study aligns with prior studies that showed HFD-induced obesity amplifies airway inflammation, with both eosinophilic and neutrophilic inflammation(9), and excess body weight in adulthood contributes to incident asthma risk in women but not men(16). Excess body fat contributes to systemic, low-grade inflammation that can impair lung function, including significant reductions in forced expiratory volume in one second (FEV_1_) and the FEV_1_/FVC ratio in obese individuals when compared to non-obese individuals(2). Further, patients with asthma undergoing bariatric surgery or diet-induced weight loss had improved pulmonary function and asthma symptom control compared to obese patients with asthma(6, 17, 18), highlighting the importance of addressing obesity in asthma management. Moreover, our results indicate that obese female patients with asthma possess a significantly higher prevalence of Th17 cells compared to their male counterparts. Prior studies have proposed that sex hormones and corresponding immune responses differ notably between males and females(9–11), implicating potential personalized therapeutic implications for managing obesity-related asthma.

Our findings also prompt future studies that explore how obesity and changes in sex hormones, like menopause, may modify asthma incidence, prevalence, and airway inflammation endotypes. During menopause, increased use of hormone replacement therapies, including estrogens, occur. Menopause represents a period of heightened asthma incidence in females(19). Our findings indicate that obesity plus estrogen signaling could promote increased Th17-mediated airway inflammation and increased neutrophil infiltration into the airway for obese females with asthma. Such endotypes of asthma are often resistant to available therapies and common among severe asthma phenotypes. Therefore, future studies exploring how hormone replacement therapy affects CD4^+^ T cell and other immune cell responses in non-obese and obese women with asthma would provide critical insight on how hormonal status and supplementation differentially affects women with asthma and contributes to treatment refractory disease.

There are limitations to our study. This study utilizes a HFD on mice that mimics but does not fully recapitulate the westernized diet. Additionally, the *Esr1*^fl/fl^ *Cd4*^Cre+^ mice will have deletion of *Esr1* in both CD4^+^ and CD8^+^ T cells, but our prior studies have shown that HDM-induced airway inflammation does not cause robust CD8^+^ T cell responses(20, 21). Therefore, eliminating ER-α signaling from CD8^+^ T cells should have minimal to no impact on the HDM-induced airway inflammation in our mouse model. While our study emphasizes the importance of ER-α signaling, it does not account for the multifaceted nature of obesity, including estrogenic potential of adipose tissue, dysbiosis of the gut microbiome, and activation of inflammatory macrophages. While these limitations are noted, our findings are still important in showing that ER-α in CD4^+^ T cells significantly contributes to exacerbated Th17 inflammation in obese female mice. Additionally, these findings underscore the necessity to consider obesity’s impact within the broader context of asthma management, particularly emphasizing hormonal factors that may inform tailored therapeutic strategies for obese women and men with asthma.

## Acknowledgements

We thank the members of the Cahill laboratories for processing clinical patients’ PBMC samples and for the GATA3 patients for participating and donating the samples. We acknowledge BioRender.com for making Figure 4A. This work was funded by the following grants from the NIH: T32 AI138932 (EHP, JCR), T32 GM139800 (EHP), R01 HL136664 (DCN, JCR), R01 HL122554 (DCN), U01AI155299 (KNC),, T32 GM007347 (KEM), and F30 HL177852 (KEM).

## Notes

### Competing Interest Statement

The authors have declared no competing interest.

