## Supplemental Figures and Tables for "Ovarian Hormones and Obesity Drive Th17-mediated Airway Inflammation through Estrogen Receptor Signaling"

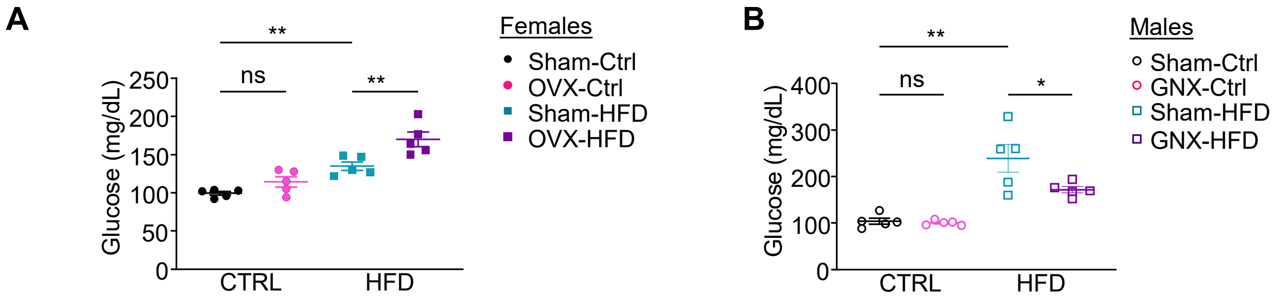


**Supplementary Figure 1: Fasting serum glucose levels in female sham-operated, female OVX, male sham-operated, or male GNX mice on control or HFD. Related to Figures 1-2.** Fasting glucose levels were determined on female sham-operated, female OVX, male sham-operated, or male GNX mice after being on control or HFD diet for 12 week. A) Female serum glucose levels. B) Male serum glucose levels. Data are expressed as mean ± SEM; n = 3-5 mice per group, $*p < 0.05$, $**p < 0.01$ one-way ANOVA with Tukey's post-hoc test.

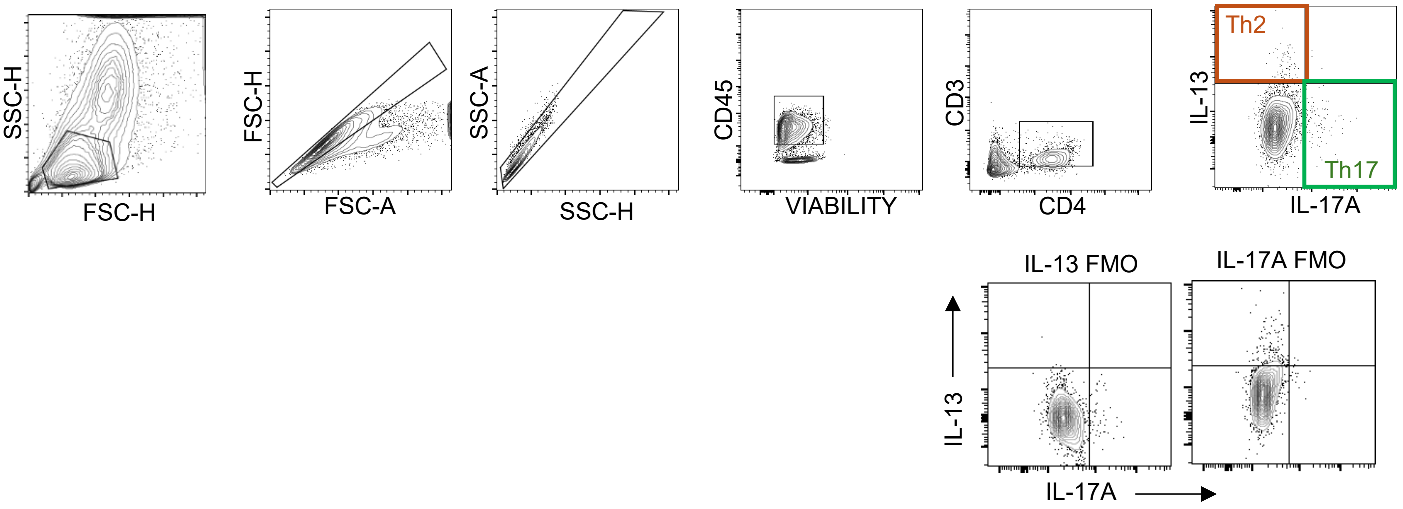


**Supplementary Figure 2: Gating strategy for identification of Th2 and Th17 cells in HDM-challenged mice.** Related to Figures 2-3.

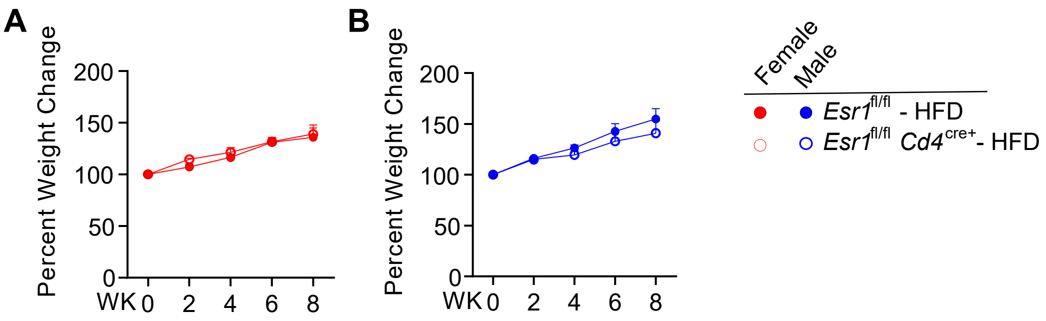


**Supplementary Figure 3: Percent weight change of *Esr1*$^{fl/fl}$ and *Esr1*$^{fl/fl}$ *Cd4*$^{cre+}$ female and male mice placed on HFD. Related to Figure 3.** Percent change in body weight from baseline in female **(A)** and male **(B)** mice fed a HFD 8 weeks.

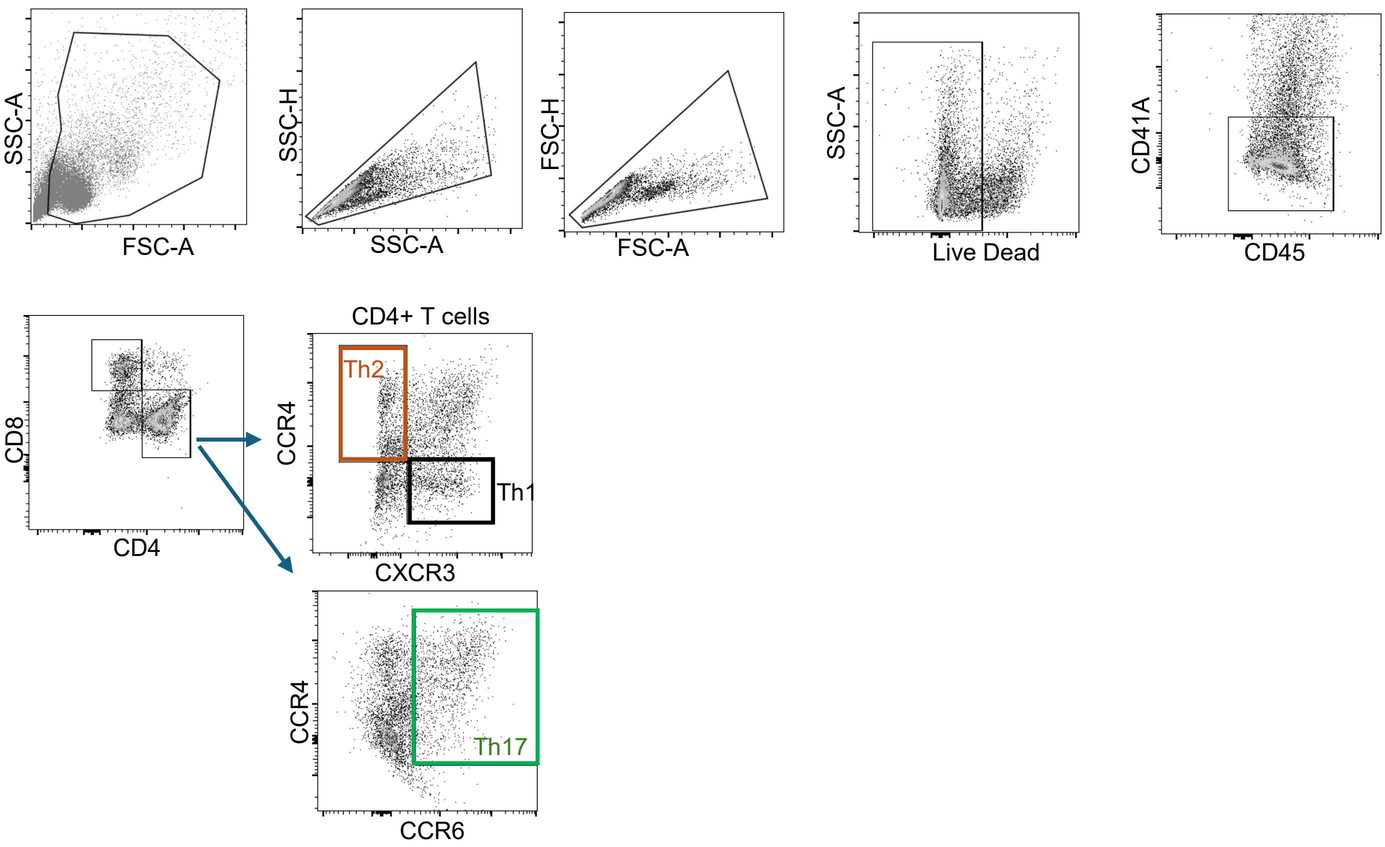


**Supplementary figure 4: Gating strategy for identification of Th1, Th2, and Th17 cells in obese women and men with asthma.** Related to Figure 4.

**Supplementary Table 1: Antibodies used in this project**.

| Antibody | Manufacturer | Catalog Number |
|---|---|---|
| Anti-Mouse CD3 | ThermoFisher Scientific | Cat #: 16-0031-82 |
| Anti-Mouse CD4 | BioLegend | Cat #: 130310 |
| Anti-Mouse CD45 | BioLegend | Cat #: 103101 |
| Anti-Mouse IL-17A | BioLegend | Cat #: 506903 |
| Anti-Mouse IL13 | ThermoFisher Scientific | Cat #: 12-7133-41 |
| Anti-Mouse IFN-γ | ThermoFisher Scientific | Cat #: 14-7311-81 |
| Anti-Mouse FOXP3 | ThermoFisher Scientific | Cat #: 14-5773-82 |
| Anti-Human CD4 | ThermoFisher Scientific | Cat #: 14-0048-82 |
| Anti-Human CD8a | BioLegend | Cat #: 301002 |
| Anti-Human CD11B | ThermoFisher Scientific | Cat #: 12-0118-42 |
| Anti-Human CD45 | ThermoFisher Scientific | Cat #: 11-9459-42 |
| Anti-Human FOXP3 | ThermoFisher Scientific | Cat #: 14-4776-82 |
| Anti-Human CD41A | ThermoFisher Scientific | Cat #: 12-0419-42 |
| Anti-Human CXCR3 | BioLegend | Cat #: 353705 |
| Anti-Human CCR4 | Miltenyi | Cat #: 130-118-496 |
| Anti-Human CCR6 | BD Biosciences | Cat #: 560619 |

**Supplementary Table 2: Clinical and Demographic Characteristics of Female and Male Obese Patients with Asthma. Related to Figure 4.**

| Characteristics | Female | Male | Combined |
|---|---|---|---|
| **N** | 6 | 4 | 10 |
| **Age (y), mean (range)** | 38.75±7 (30-50) | 39.25±7 (30-47) | 39±7 (30-50) |
| **Race, White/Black/Asian/Other** | 3/1/1/1 | 4/0/0/0 | 7/1/1/1 |
| **Ethnicity, Non-Hispanic/ Hispanic** | 5/1 | 4/0 | 9/1 |
| **Age(y) of asthma onset, mean (range)** | 29.2±13.42 (1-38) | 13.5±11 (1-26) | 22.9±14.6 (1-38) |
| **BMI ($kg/m^2$), mean (range)** | 35.6±5 (29.4-41.6) | 35.5±7 (28.1-46.8) | 35.6±6 (28.1-46.8) |
| **Mean ACQ score ± SD** | 2.95±0.39 | 2.45±0.5 | 2.75±0.5 |
